# Automated home-cage sipper devices reveal age and sex differences in ethanol consumption patterns

**DOI:** 10.1101/2023.03.22.533844

**Authors:** Rachel C. Rice, Annalisa M. Baratta, Sean P. Farris

## Abstract

**Background:** Alcohol consumption and alcohol use disorder (AUD) are influenced by circadian rhythms. Alcohol-related behaviors and circadian rhythms differ with age and sex, but little preclinical research has compared circadian patterns of ethanol consumption across ages and sexes. Cost-effective tools are becoming widely available that could elucidate these patterns, including open-source, Arduino-based home-cage sipper devices. We hypothesized these devices would reveal age- and sex-specific patterns of ethanol and water consumption and ethanol-induced changes in overall fluid consumption rhythms.

**Methods:** We used Arduino-based home-cage sipper devices in a continuous-access two-bottle choice (2BC) paradigm with water and ethanol (10% v/v) for 14 days to measure drinking patterns of male and female adolescent (3-week), young adult (6-week), and mature adult (18-week) C57BL/6J mice. Grams per kilogram (g/kg) fluid consumption were manually recorded at the beginning of each dark cycle, while sipper devices continuously recorded drinking.

**Results:** Manually collected 2BC data confirmed greater ethanol consumption in females of all age groups as well as stepwise decreases in total fluid consumption with age in ethanol-exposed females and in both sexes in the water-only groups. Correlations of manually recorded versus sipper count data were strong and significant for ethanol, but modest for water, suggesting fluid differences in drinking microstructure. Blood ethanol concentrations were significantly correlated with sipper data, indicating sipper devices accurately determine temporal ethanol consumption. Sipper devices uncovered subtle differences in circadian rhythms by age, group, and ethanol exposure, including minor phase advancement in ethanol consumption with age in females, but delay in males, as well as a female-specific phase delay and weaker amplitude in total fluid consumption across age groups with ethanol exposure.

**Conclusions:** We show these devices capture circadian patterns of ethanol drinking behavior that differ by age, sex, and ethanol exposure inaccessible by manual 2BC measurements alone. Future studies may examine older age groups and further dissect circadian drinking patterns by age and sex and their molecular mechanisms relevant to AUD pathogenesis.

## Introduction

Alcohol use disorder (AUD) impacts ∼28 million individuals ages 12 and older in the United States, with an estimated lifetime prevalence of 36% in males and 22.7% in females. (Grant et al., 2015, Flores-Bonilla and Richardson, 2020, Substance Abuse and Mental Health Services Administration (SAMHSA), 2024) Alcohol consumption and AUD are influenced by age (Han et al., 2017, Substance Abuse and Mental Health Services Administration (SAMHSA), 2024, White et al., 2023) and sex, (Han et al., 2017, Substance Abuse and Mental Health Services Administration (SAMHSA), 2024, White et al., 2023, Agabio et al., 2017, White, 2020) and human and preclinical studies have demonstrated sex differences in alcohol consumption emerge during adolescence. (White, 2020, Truxell et al., 2007, Witt, 2007)

A bidirectional relationship exists between ethanol exposure and circadian rhythms, which are also influenced by age and sex. Circadian rhythms weaken with age in humans and animal models, with sleep/wake, core body temperature, hormone secretion, locomotor activity, and eating patterns displaying decreased amplitude. (Banks et al., 2016, Nakamura et al., 2016, Nakamura et al., 2015, Duffy et al., 2015) In humans, sleep timing displays phase delay during puberty, peaking at ∼age 20, then increasingly advances with age. (Roenneberg et al., 2004, Banks et al., 2016, Nakamura et al., 2016) Sex appears to interact with age to influence circadian rhythms, but results differ across studies and measures analyzed. (Carrier et al., 2017)

Acute ethanol exposure disrupts sleep, and individuals with AUD are more likely to report sleep disturbances. (Hasler et al., 2015) Conversely, disruption in circadian rhythms, such as in shift work and jet lag, increases risk for excessive drinking and AUD. (Hasler et al., 2015) Rodent studies have noted increased sensitivity to circadian rhythm disruption with ethanol exposure in adults compared to adolescents. (Ruby et al., 2017) Although preclinical studies have begun to dissect the interaction of age and sex on circadian rhythms (Bellfy et al., 2025, Chang et al., 2026, Ruby et al., 2017) and ethanol consumption, (Foo et al., 2023, Piekarski et al., 2022, Waldron et al., 2024, Collins et al., 1975, Melon et al., 2013, Mackowiak et al., 2025, Cuitavi et al., 2024, Ruby et al., 2017, Garcia-Burgos et al., 2009) a dearth of research directly compares circadian patterns of voluntary alcohol consumption or ethanol exposure-related disruption in circadian rhythms across age and sex in wild-type animals.

The current study utilizes cost-effective, open-source automated home cage sipper devices adapted from Godynyuk *et al*. (2019) (Godynyuk et al., 2019) in a continuous access two-bottle choice (2BC) paradigm between 10% ethanol and water to examine age- and sex-related differences in ethanol consumption in wild-type C57BL/6J mice. Water consumption was analyzed in controls to determine whether circadian patterns of fluid consumption are disrupted by age and sex with ethanol exposure. We hypothesized these sipper devices would reveal age- and sex-specific patterns of ethanol and water consumption not accessible by manual 2BC measurements, as well as changes in overall fluid consumption rhythms with ethanol exposure that differ by age and sex.

Our findings validate these sipper devices in measuring temporal patterns of ethanol consumption behavior. Sipper devices revealed subtle changes in temporal drinking patterns by age, sex, and ethanol exposure not detectable by manual measurement, as well as predicted blood ethanol concentrations at projected peak ethanol consumption. Taken together, our findings profile age- and sex-related circadian patterns of ethanol consumption behavior in C57BL/6J mice that can be expanded upon by further research.

## Methods

### Animals

Overall experimental design is summarized in **Figure 1A**. Male and female (n=36/sex) C57BL/6J mice were purchased from the Jackson Laboratory (Strain #000664; Bar Harbor, ME, USA). Age groups consisted of mice 3-, 6-, and 18 weeks of age (n=12/age/sex) at the time of receipt. These ages were chosen to represent adolescent, young adult, and mature adult mice, respectively, following guidelines supported by The Jackson Laboratory. (Dutta and Sengupta, 2016, Flurkey et al., 2007, 2017) Mice were habituated to the University of Pittsburgh animal facility for 48h, then singly-housed on ventilated racks in a temperature- and humidity-controlled room with *ad libitum* access to food and water under 12/12h reverse light-dark cycle (lights on at 22:00, off at 10:00). Mice were habituated to the reverse light-dark cycle for at least five days before experiments began. All experiments were approved by the Institutional Animal Care and Use Committee of the University of Pittsburgh and conducted in accordance with the National Institutes of Health Guidelines for the Care and Use of Laboratory Animals.

**Figure 1.**
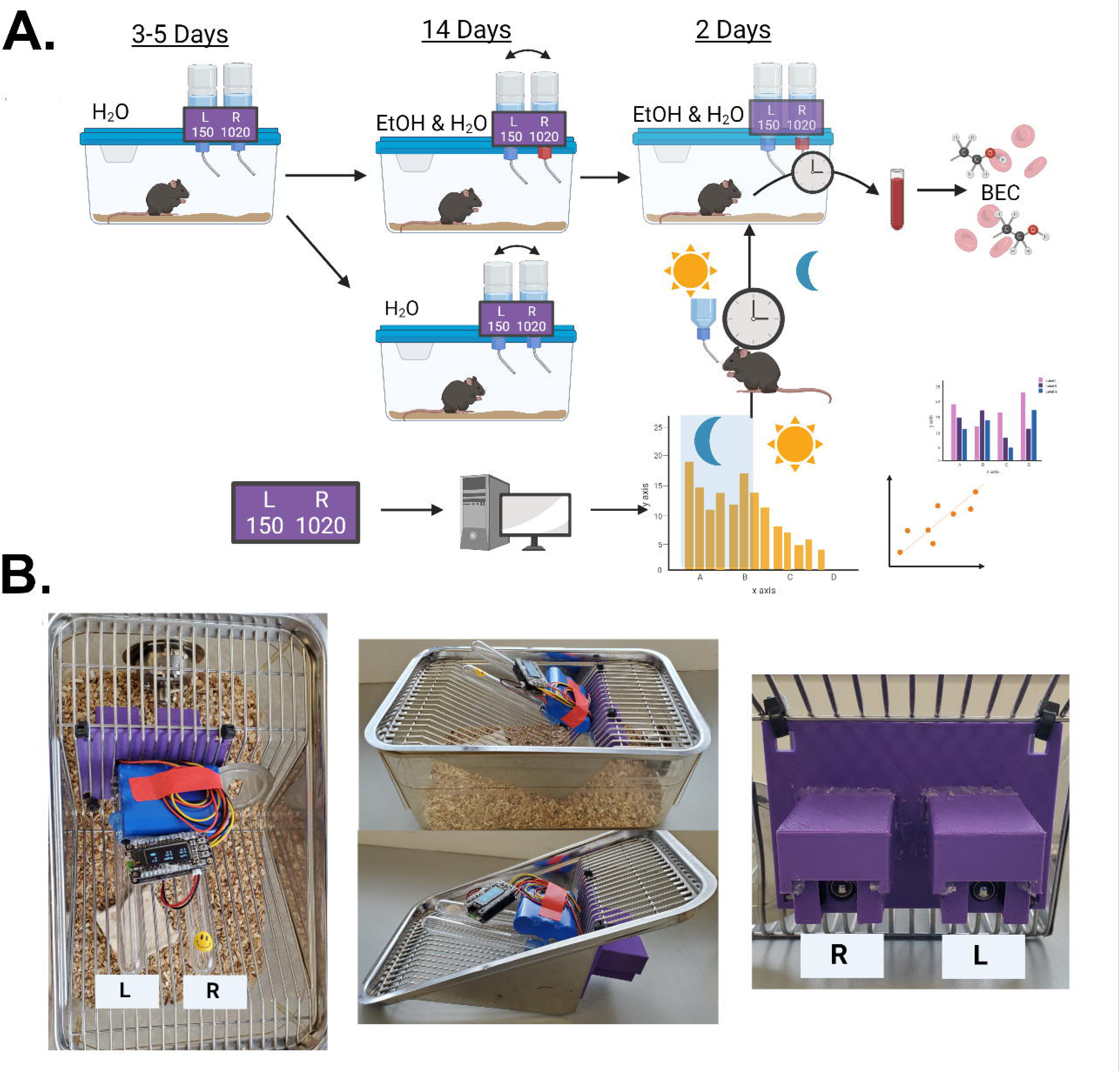
Study overview and sipper devices. (A.) Experimental overview and timeline. Numbers below “L” and “R” are representative of sipper counts. (B.) Photographs of sipper device two-bottle choice setup on an empty mouse cage. Figure 1A *created using Biorender.com*.

### Construction of home-cage sipper devices

Automated home-cage sipper devices were constructed following Godynyuk *et al*. (2019) (Godynyuk et al., 2019) with minor modifications. A new device housing was constructed to allow for the use of Amuza ball bearing sipper tubes (Drinko Measurer Sipper Tube, 3.5in) within 10-mL glass bottles (Amuza, San Diego, CA) that allow for easy daily bottle refilling and switching. The housing was designed using Tinkercad and printed on a Sindoh 3DWOX1 printer using polylactide thermoplastic. STL files for the custom sipper housing are available at: https://github.com/rachelcrice/Homecage_Sippers.

Devices were secured on the wire top of each cage using plastic cable ties and were positioned to ensure mice maintained sufficient access to food, which was distributed across the cage top equally near both bottles to avoid food-related side preference. Two sipper tubes were placed into the sipper apparatus. **Figure 1B** illustrates the completed sipper apparatus in an empty cage. An earlier version of the device Python code by Godynyuk *et al*. (dated October 6, 2018) was used that collected data every ten seconds.

### Continuous access two-bottle choice

Mice were acclimated to the sipper devices for 3-5 days with water before continuous access 2BC began. 2BC was performed for 14 days. Half of each group (n=6/age/sex) was given a choice between ethanol (10%; v/v) and water, while the other half received water only. Every 24h, all bottles were weighed and switched in position and data from the devices were collected by experimenters. Bottles were placed into the device apparatus such that each sipper was flush with the plastic housing, meaning mice could equally access each sipper and the sippers did not interfere with the device photobeams. An empty control cage with the device housing and sipper tubes was maintained to account for spillage. During experiments, left and right side were defined as experimenter left/right facing the cage front (**Figure 1B**).

Peak ethanol consumption period was determined via circadian analysis of daily sipper device data within subjects (see below). For two days following 2BC, the ethanol bottle was maintained on the left side and tail vein blood was collected from ethanol mice at approximate predicted peak of ethanol consumption. Water-only mice also underwent blood collection, but this blood was not analyzed. Blood was centrifuged for two minutes at 12,000 rpm in heparin-coated glass capillary hematocrit tubes (Fisher Scientific, Cat #22362566) in a Unico Microhematocrit centrifuge (Model C-MH30, Dayton, NJ). Serum analyzed for ethanol concentration on an Analox Alcohol Analyzer (AM1, Analox Instruments, London, UK). Blood ethanol concentration (BEC) measurements were corrected using a standard curve constructed via a serial dilution of Analox 100 mg/dL standards in distilled water.

### Statistical analyses of manually-collected 2BC data and end-day sipper counts

All statistical analyses were performed in the R statistical environment (v.4.5.0). (Team, 2025) Ethanol, water, and total fluid consumption were calculated in grams per kilogram (g/kg) body weight for each day from bottle weight data and analyzed by day as well as summed across the 14-day period. Ethanol preference was calculated as the percent grams of fluid consumed from the ethanol bottle over total gram consumption from bottle weights. Side preference in the water group was calculated as the percent grams of fluid consumed from the right bottle over total gram consumption. Daily g/kg consumption data were analyzed across Sex, Age, and Day via a linear mixed effects model using the lmerTest (v.3.1.3) R package. (Kuznetsova et al., 2017) Total g/kg consumption, average ethanol preference, and average side preference were analyzed across Sex and Age via 2×3 ANOVAs using the stats package in base R. (Team, 2025) Statistically significant interaction effects were analyzed via Tukey post-hoc tests using the emmeans R package. (Lenth R, 2026) Linear regression analyses were performed on g/kg ethanol or water consumption from experimenter measurements as well as BECs versus end-day sipper device counts. Measurements with null sipper values due to loss of battery power or other device errors were omitted from analysis. Pearson correlation coefficients were calculated using the stat_cor function from the ggpubr R package (v.0.6.3). (Kassambara, 2026) Terms such as “strong” or “weak” to denote correlation strength were utilized according to previously-discussed guidelines for biomedical research to discuss correlation coefficients. (Schober et al., 2018, Akoglu, 2018)

### Circadian analysis of sipper counts

A custom R pipeline was utilized for circadian analysis of sipper count data. Raw cumulative counts recorded every 10 seconds were differenced to yield interval counts, then binned into 1-minute and subsequently 60-minute intervals. Ethanol and water counts were assigned to the correct bottle based on the recorded side position for each recording day. Mean 60-minute counts were averaged per animal across the 14-day recording period. A linear cosinor model was fit to each animal’s mean 60-minute binned counts using ordinary least squares: *y* = *MESOR*+ *βcos*(*ωt*) + *γsin*(*ωt*), where 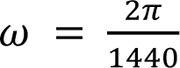, corresponding to a 24-hour period in minutes, and MESOR refers to the midline estimating statistic of rhythm, the rhythm-adjusted 24-hour mean sipper counts. MESOR, amplitude (√*β*^2^ + *γ*^2^, half the peak-to-nadir range), and acrophase (approximate peak, determined by evaluating the fitted curve at each minute across the 24-hour window and identifying the maximum) were extracted for each animal. These parameters were averaged within Age × Sex groups to produce average cosinor curves. Group differences in MESOR, amplitude, and acrophase were assessed using two-way ANOVA (Age × Sex), with post-hoc Tukey’s HSD where statistically significant interactions were detected. All data and analysis code are available at: https://github.com/rachelcrice/Homecage_Sippers.

## Results

### Manual analysis of ethanol & water consumption in a continuous access 2BC paradigm

To determine the effects of age and sex on ethanol, water, and total fluid consumption, male and female mice, aged 3-, 6-, and 18-weeks, were allowed free access to one bottle containing ethanol and one containing drinking water for 14 days. A separate group comprised water-only controls. Female animals consumed between approximately 5-15 g/kg ethanol per day, while male animals consumed ∼2.5-10 g/kg ethanol per day. Three-way repeated measures ANOVA of daily ethanol intake revealed main effects of Sex and Day (p < 0.01), but no main effect of Age nor interaction effects (**Figure 2A**). Two-way ANOVA on total g/kg ethanol consumption over the 14-day period similarly revealed a main effect of Sex (p < 0.01), but no main effect of Age (**Figure 2B**). Although this effect was not statistically significant, female 3-week mice drank more ethanol than 6- or 18-week mice, which displayed similar average patterns of daily and cumulative ethanol consumption; male mice showed similar patterns of daily and cumulative ethanol consumption across groups (**Figure 2A-B**). There were no statistically significant effects on daily ethanol preference (**Figure 2C**), though two-way ANOVA on average ethanol preference also yielded a main effect of Sex (p < 0.01) (**Figure 2D**).

**Figure 2.**
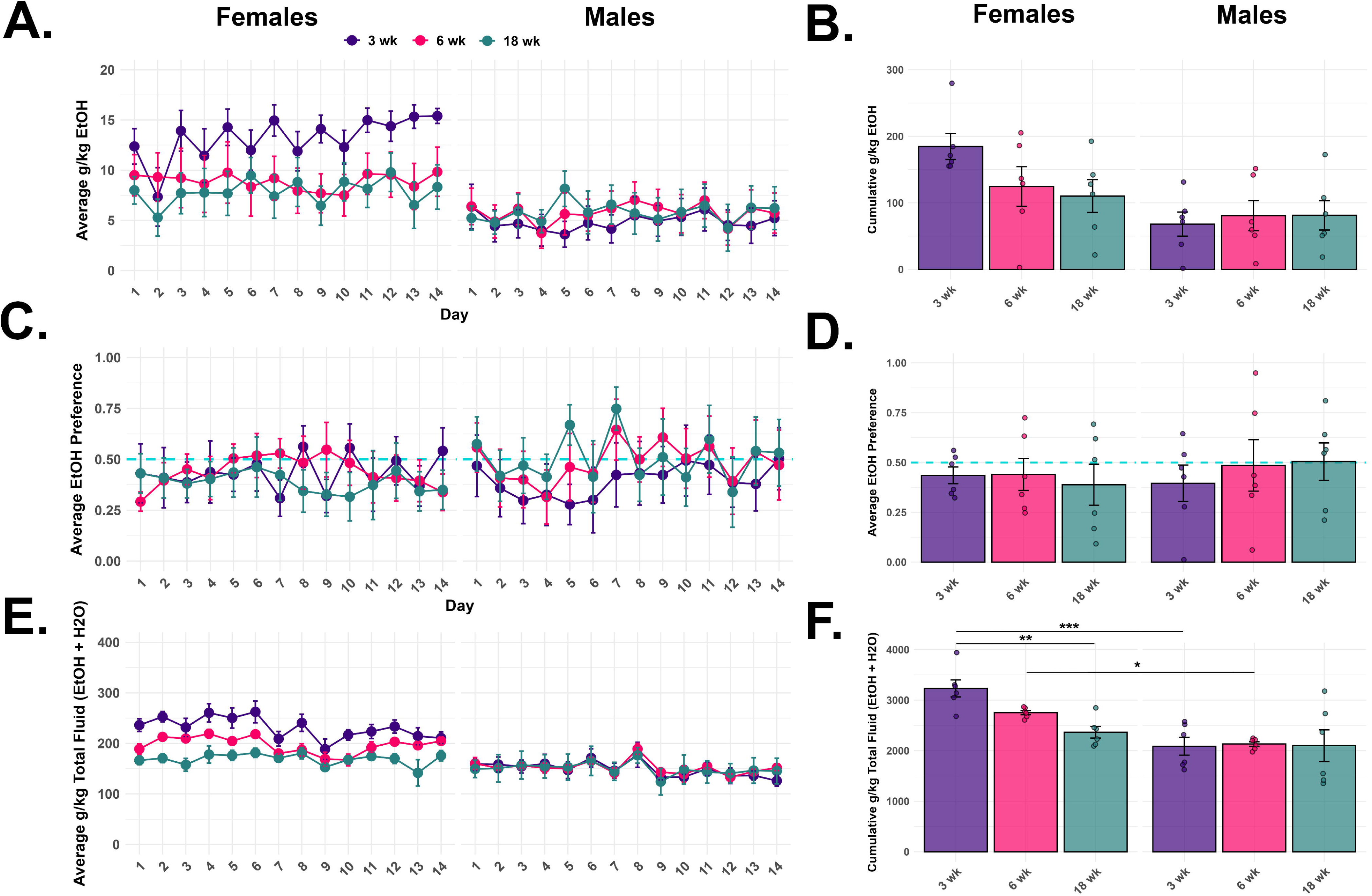
Manually recorded two-bottle choice data from the ethanol-exposed group by age and sex. (A.) Average daily grams per kilogram (g/kg) ethanol consumption. (B.) Cumulative g/kg ethanol consumption. (C.) Average daily ethanol preference. (D.) Average ethanol preference over the 14-day period. (E.) Average daily total g/kg fluid (ethanol + water) consumption. (F.) Cumulative total g/kg fluid consumption.

Total g/kg fluid consumption was summed between the ethanol and water bottles to determine overall consumption behavior in ethanol group mice (**Figure 2E**). Daily total fluid consumption in the ethanol-exposed group displayed main effects of Sex and Day (p < 0.001) as well as Age x Sex (p < 0.05) and Day x Sex (p < 0.001) interaction effects; the same was seen in total fluid consumption summed over the 14-day access period. In both analyses, Age approached significance (p = 0.058; **Figure 2F**), and Tukey post-hoc analyses found statistically significant differences between sexes in the 3-week (p < 0.001) and 6-week (p < 0.05) age groups, as well as a statistically significant difference between the 3-week and 18-week age groups in females (p < 0.01; **Figure 2F**).

Total water consumption was assessed in the water-only group by summing daily g/kg consumption from both water bottles. This revealed statistically significant main effects of Age, Sex, and Day (p < 0.001) as well as a Sex x Day interaction (**Figure 3A**). The main effects of Age and Sex were preserved in cumulative water consumption (p < 0.001; **Figure 3B**). Male and female mice of all age groups preferred the left bottle in the absence of ethanol; this effect was stronger in the males (main effect of Sex; p < 0.01; **Figure 3C**).

**Figure 3.**
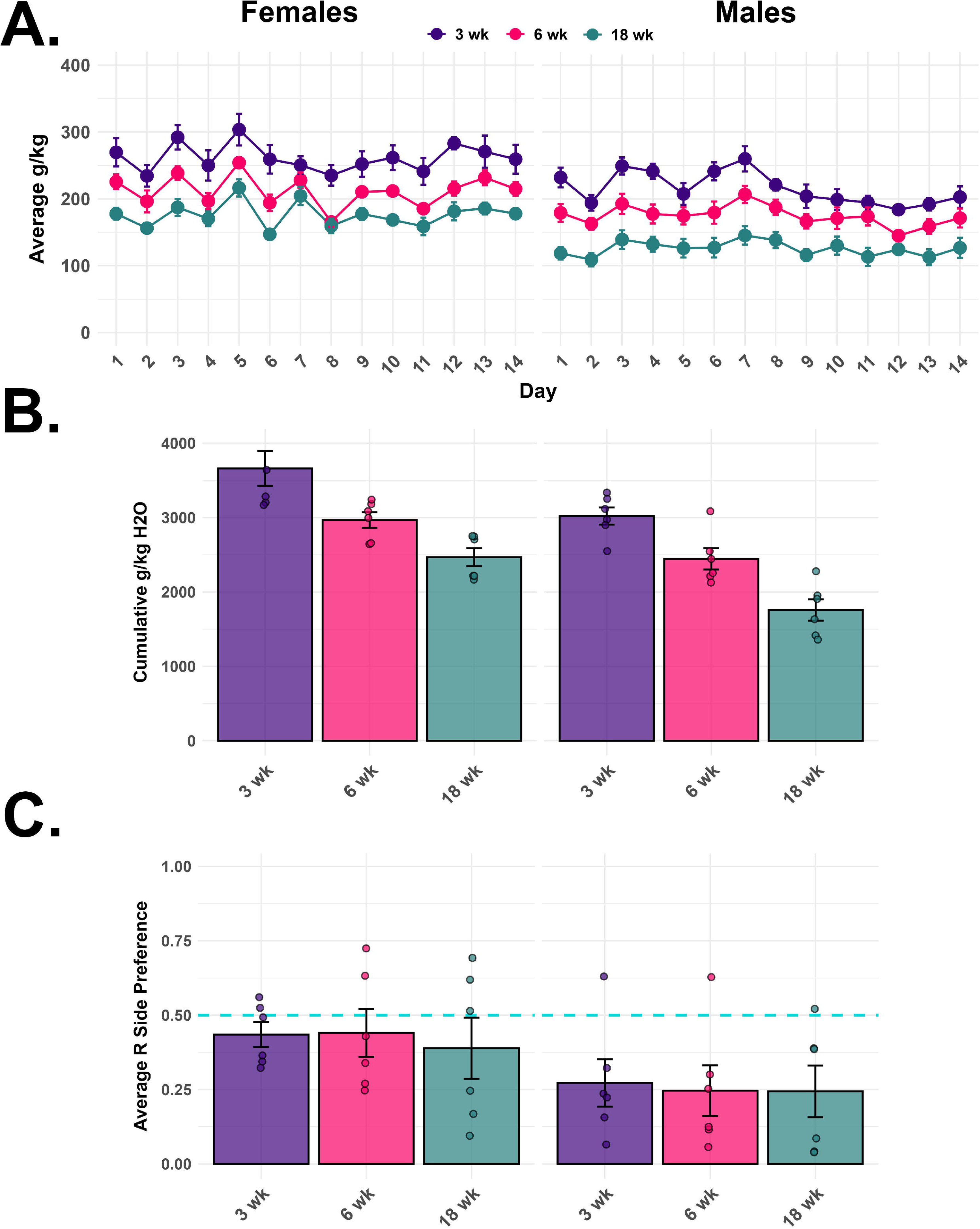
Manually recorded two-bottle choice data from the water-only group by age and sex. (A.) Average daily grams per kilogram (g/kg) water consumption. (B.) Cumulative g/kg water consumption. (C.) Average right side preference.

### Correlation analysis of manual and sipper device 2BC data

Linear regressions were performed to determine the accuracy of the sipper devices in measuring the amount of ethanol and water consumed by mice. Correlations were first performed within Sex, collapsed across Age to increase statistical power. Device counts showed strong, statistically significant correlations with manual 2BC data for ethanol consumption in males and females (Pearson R = 0.82-0.83, p < 0.001; **Figure 4A**). Total fluid consumption in the ethanol group had weaker correlations with summed sipper counts; for females, the correlation was not statistically significant, and for males, a weak significant correlation was observed (R = 0.36, p < 0.001; **Figure 4B**). Similarly, in the water-only group, sipper counts displayed weak-to-moderate, statistically significant correlations with manual 2BC data (Pearson R = 0.33-0.4, p < 0.001; **Figure 4B**). Given stratification of scatter plots by age as well as differential water consumption among age groups in the water-only group, correlation analyses were repeated within group and sex. For ethanol consumption, correlations remained statistically significant and strong for all Sex x Age groups (R > 0.70, p < 0.001; **Supplementary Figure S1**). For total fluid consumption in the ethanol group, correlations were statistically significant, but weak for 3-week males and females (R < 0.04, p < 0.01) and significant, but of moderate strength for 18-week males (R = 0.61; **Supplementary Figure S2**). For total water consumption in the water-only group, Pearson correlations were statistically significant except for in the 18-week females, though all significant correlations were of weak-to-moderate strength (R = 0.26-0.57, p < 0.05; **Supplementary Figure S3**).

**Figure 4.**
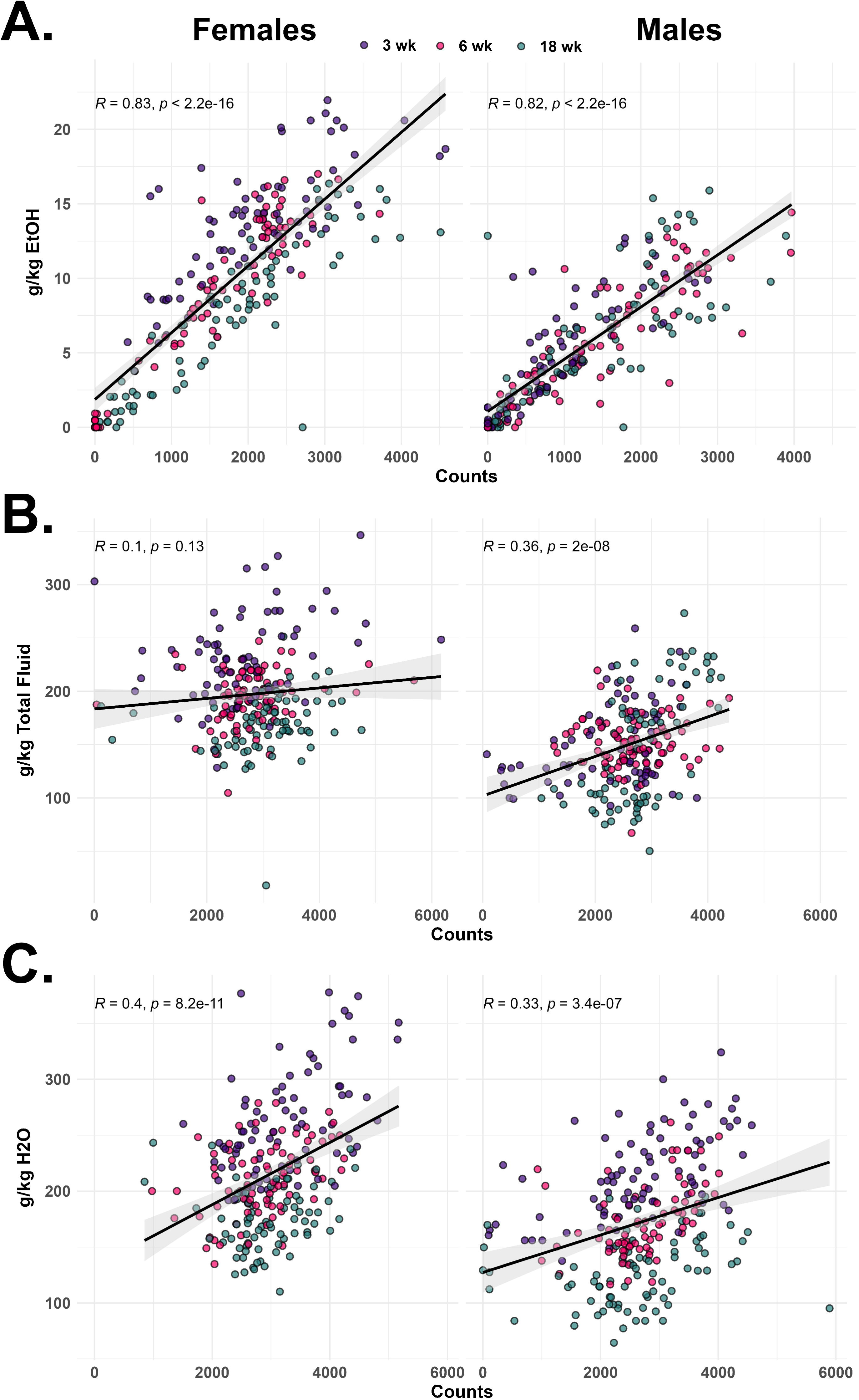
Correlations of manually collected g/kg two bottle choice data versus sipper device counts by sex for: (A.) Ethanol consumption in the ethanol-exposed group; (B.) Total fluid consumption (ethanol + water) in the ethanol-exposed group; (C.) Water consumption in the water-only group.

To determine whether circadian sipper data could predict BECs at approximate peak ethanol consumption period, sipper counts at acrophase predicted from individual subjects’ 30-minute binned sipper data were correlated to the BECs measured at this time for the ethanol group. This correlation was statistically significant, but of weak-to-moderate strength (**Figure 5**).

**Figure 5.**
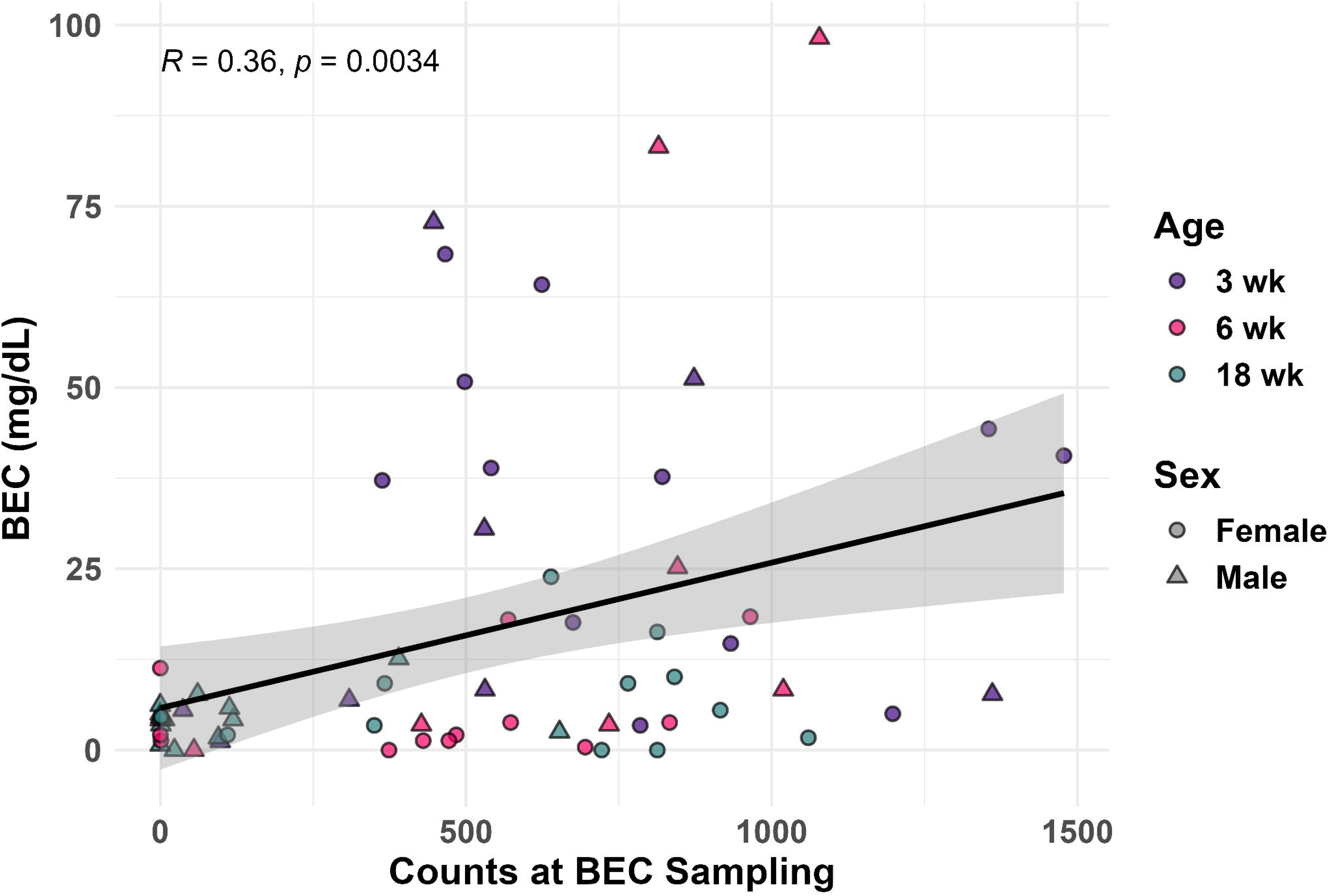
Correlations of blood ethanol concentrations (BECs) in milligrams per deciliter (mg/dL) versus sipper counts at the time of BEC sampling, which was performed at predicted peak ethanol consumption from individual circadian sipper device data.

### Circadian differences in ethanol and water consumption by age and sex

Sipper device count data were analyzed in R in 60-minute bins to construct circadian histograms averaging 14-day fluid consumption within groups. For the ethanol group, both ethanol and total (*i.e.,* ethanol + water) counts were analyzed (**Figure 6A-B**). Total water counts across the two bottles were analyzed for the water-only group (**Figure 7**). Across groups, greater counts were observed during the dark cycle than the light cycle (**Figures 6-7**). To quantitatively analyze circadian patterns of consumption, 60-minute binned data underwent cosinor modeling (**Figure 8**). Circadian cosine curves for each age/sex group for each 2BC measure were generated (**Figure 8A, C, E**), and acrophase, amplitude, and MESOR were calculated for each age/sex group for ethanol and total fluid counts (ethanol group) as well as total water counts (ethanol naïve group; **Figure 8B, D, E**). For ethanol consumption acrophase, there was an Age x Sex interaction effect (p < 0.01). Post-hoc testing did not reveal any statistically significant differences when adjusted for multiple comparisons, though comparisons between 18-week and 3-week females as well as 18-week males and females approached significance (p_adj_ = 0.051 and 0.057, respectively; **Figure 8A-B**). However, qualitative comparisons of ethanol consumption acrophase still suggest potentially biologically meaningful differences in peak ethanol consumption across age groups, namely subtle phase advancement in females, but delay in males with age (**Figure 8A-B**). Ethanol consumption acrophase was approximately 6 hours into the dark phase for 3-week female mice and approximately 5 hours into the dark phase for their male counterparts. For 6-week mice, ethanol consumption acrophase was approximately 5 hours for females and 4.5-5 hours for males. In the 18-week mice, ethanol consumption acrophase was around 4 hours into the dark cycle for females and 5.5-6 hours for males (**Figure 8A-B**). Ethanol count amplitude and MESOR did not display any statistically significant differences across sex/age groups, though average MESOR was approximately halved in 18-week males compared to females (**Figure 8B**).

**Figure 6.**
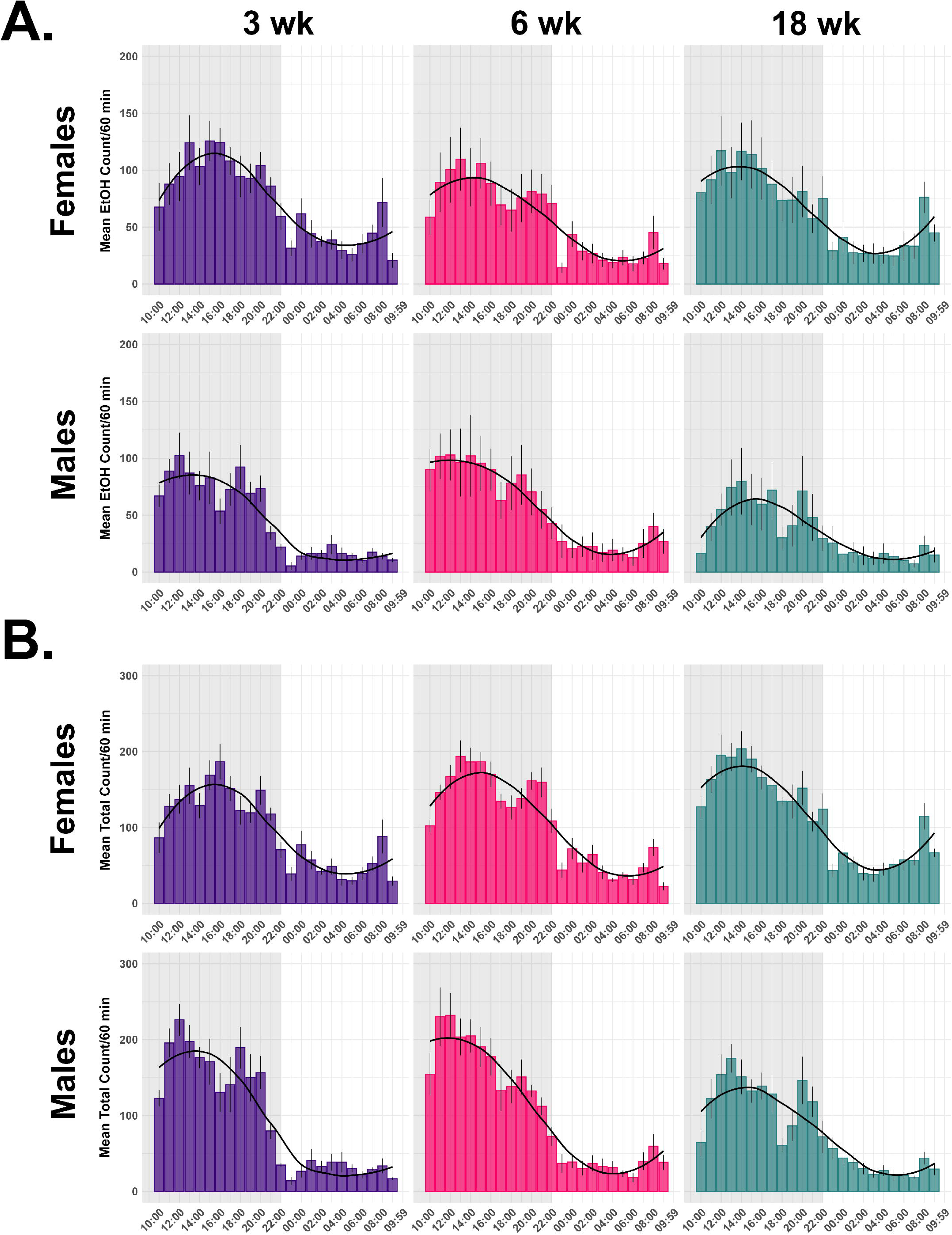
Sipper device data in 60-minute bins averaged within age-sex groups for the ethanol-exposed group. (A.) Ethanol counts; (B.) Total (ethanol + water) counts.

**Figure 7.**
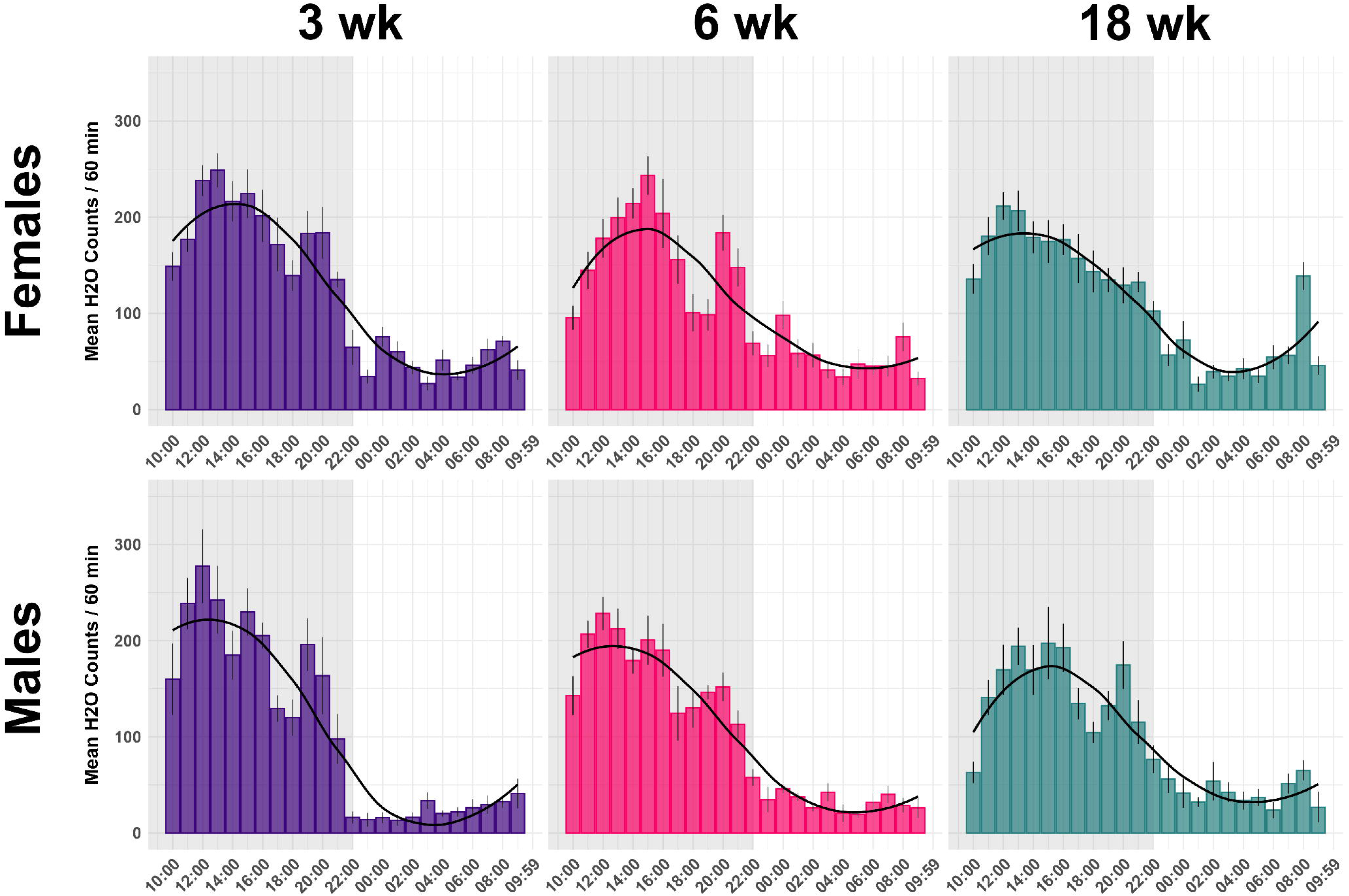
Sipper device data in 60-minute bins averaged within age-sex groups for the water-only group.

**Figure 8.**
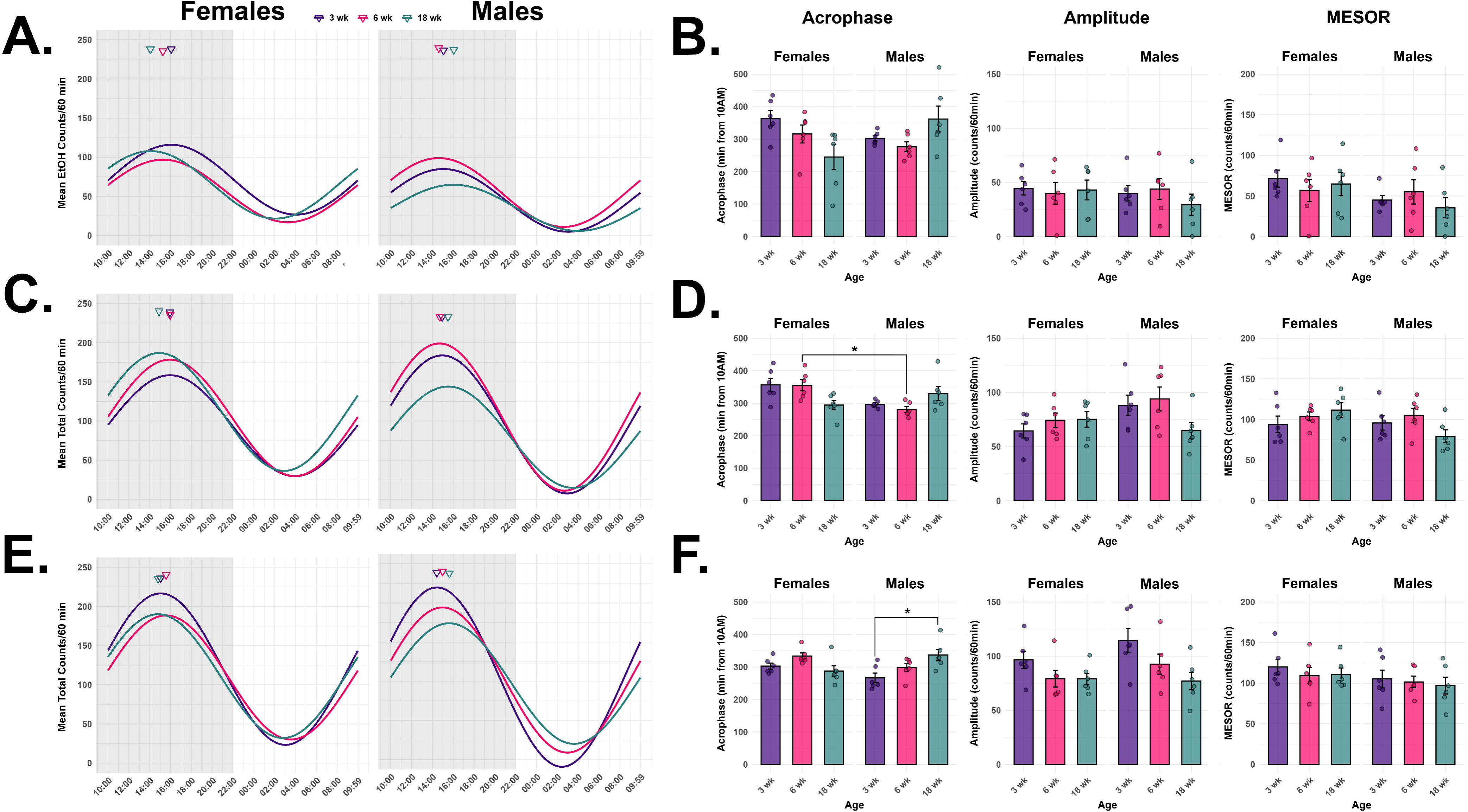
Cosinor circadian analysis of sipper device data in 60-minute bins. (A.) Average cosine curves by sex and age group for sipper device ethanol counts from the ethanol-exposed group. (B.) Acrophase, amplitude, and midline estimating statistic of rhythm (MESOR) for ethanol counts in the ethanol-exposed group by age and sex. (C.) Average cosine curves by sex and age group for sipper device total (ethanol + water) counts from the ethanol-exposed group. (D.) Acrophase, amplitude, and midline MESOR for total (ethanol + water) counts in the ethanol-exposed group by age and sex. (E.) Average cosine curves by sex and age group for sipper device total water counts from the water-only group. Acrophase, amplitude, and MESOR for total water counts in the water-only group by age and sex.

Total consumption acrophase in the ethanol group displayed a main effect of Age (p < 0.05) as well as an Age x Sex interaction effect (p < 0.01; **Figure 8C-D**). The only statistically significant difference in total count acrophase when adjusted for multiple comparisons was between 6-week males and females (p_adj_ < 0.05; **Figure 8D**). Total count acrophase was approximately 5.5-6 hours into the dark cycle for 3-week females and 5 hours for males. These acrophase patterns were preserved within sexes in the 6-week group. At 18-weeks of age, total count acrophase was approximately 5 and 5.5 hours into the dark cycle for females and males, respectively (**Figure 8C-D**). Total counts did not statistically differ in amplitude nor MESOR across age/sex, though male 18-week amplitude was approximately 30 counts/hour lower than the 3-week and 6-week groups (**Figure 8D**).

In the water-only group, total water acrophase displayed a statistically significant Age x Sex interaction effect (p < 0.01), while Age approached statistical significance (p = 0.059; **Figure 8E-F**). The only statistically significant difference in total water count acrophase when adjusted for multiple comparisons was between 18-week and 3-week males (p_adj_ < 0.05). Acrophase was approximately 5-6 hours into the dark cycle for all female age groups; for males, Acrophase increased stepwise with age from approximately 4 to 6 hours into the dark cycle (**Figure 8E-F**). Amplitude displayed a main effect of Age (p < 0.01). In females, amplitude decreased from nearly 100 counts/hour in the 3-week group to ∼75 counts/hour in the 6- and 18-week groups. In males, amplitude decreased almost stepwise with age from approximately 110-75 counts/hour. MESOR did not statistically differ between groups (**Figure 8F**).

Finally, total fluid consumption cosinor parameters were compared across ethanol and water-only groups to determine whether ethanol exposure differentially affects circadian drinking rhythms across ages, within sexes. In females, acrophase displayed main effects of Age (p < 0.01) and Exposure Group (p < 0.05; **Figure 9**). Overall, acrophase decreased with age, and ethanol-exposed females showed a later acrophase (suggesting phase delay) than their water-only counterparts. Amplitude displayed a main effect of Group in females, where ethanol-exposed mice had weaker rhythms than water-only mice, though this appears to have primarily been driven by the 3-week age group (p < 0.05; **Figure 9**). In males, acrophase and amplitude had main effects of Age (p < 0.01), where acrophase increased (suggesting phase delay) and amplitude decreased with Age (suggesting weaker rhythms) across exposure groups (**Figure 9**). In both males and females, there were no statistically significant effects on MESOR (**Figure 9**).

**Figure 9.**
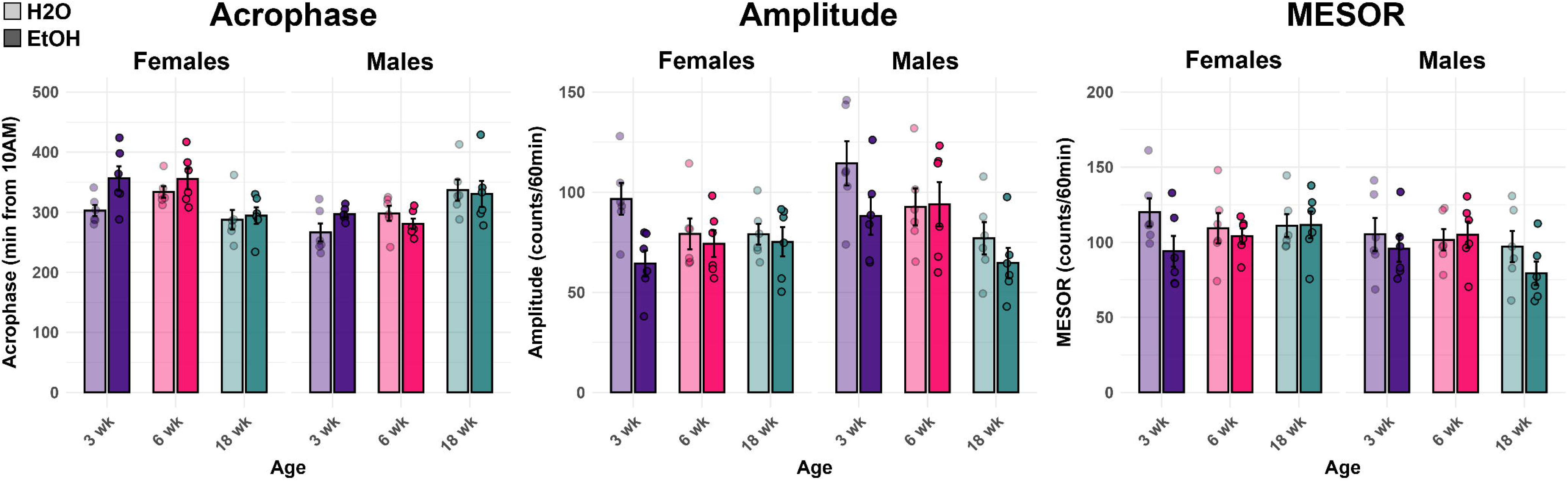
Comparison of cosinor statistics by age and sex across water-only and ethanol groups for total fluid consumption (water in the water-only group, ethanol + water in the ethanol group). Acrophase, amplitude, and midline estimating statistic of rhythm (MESOR) are given.

## Discussion

Open-source automated home-cage sipper devices adapted from Godynyuk *et al*. (2019) (Godynyuk et al., 2019) were validated in a 14-day, continuous access 2BC paradigm to investigate age- and sex differences in ethanol consumption and circadian patterns of ethanol and water consumption. Correlations of manually collected g/kg 2BC consumption data versus automated sipper counts differed in strength by age, sex, and the measure of consumption. Ethanol bottle correlations were strong and statistically significant across age and sex, while summing total consumption and sipper counts across ethanol and water bottles for the ethanol-exposed groups resulted in weaker correlations that were not statistically significant in some age and sex groups. Total water correlations in the ethanol-naïve group were the weakest, with less statistically significant prediction of fluid consumption from sipper counts. Weaker correlations for total fluid (in the ethanol-exposed group) and water (in the ethanol-naïve group) could be due to differing drinking microstructures, namely longer bouts of drinking water with fewer individual sips recorded versus shorter bouts of ethanol with more frequent sips; similarly, differences in the predictive value of sipper counts in measuring fluid consumption groups could reflect differing drinking microstructure by age and/or sex. (Godynyuk et al., 2019, Mackowiak et al., 2025, Robinson and McCool, 2015, Barkley-Levenson and Crabbe, 2012, Petersen et al., 2024, Caruso et al., 2021, Yoon et al., 2025) Our average water consumption correlations as well as variation in water consumed versus counts among individuals, groups, and days recapitulated the findings of Godynyuk *et al*. in validating these devices. (Godynyuk et al., 2019) Additionally, we found binned circadian ethanol exposure data could moderately predict blood ethanol concentrations at expected peak. This modest effect size could be attributable to the fact that mice were not voluntarily drinking to intoxication (as all BECs were below 100 mg/dL, with most falling below 50 mg/dL), as well as device errors in recording counts and other non-drinking related interruptions of the photobeams.

Experimenter-collected data revealed increased ethanol consumption in female compared to male animals in daily and cumulative intake. Although no statistically significant effects of age on ethanol consumption or preference were observed, 3-week female mice drank the most ethanol out of any age groups. When water and ethanol fluid consumption were summed in the ethanol-exposed group, females showed a stepwise decreasing pattern of overall consumption with age, where the 3-week age group consumed significantly more total fluid than the 18-week group. Except for the 18-week group, females of each age group consumed significantly more total fluid than males. Males of all age groups showed similar levels of total fluid consumption. In the ethanol-naïve group, females drank significantly more water than males, and another statistically significant stepwise decreasing effect of age was observed in total water consumption.

Higher voluntary ethanol intake in adult females in the 2BC paradigm has previously been documented in rodent alcohol research. (Salazar and Centanni, 2024, Tambour et al., 2008, Cuitavi et al., 2024) This sex difference has also been observed during rodent adolescence. (Westbrook et al., 2018, Tambour et al., 2008) Similarly, adolescent rodents have been found to have elevated voluntary ethanol consumption relative to adults. (Moore et al., 2010, Morales et al., 2014, Tambour et al., 2008, Cuitavi et al., 2024, Garcia-Burgos et al., 2009) We did not observe increased voluntary ethanol intake in adolescent males relative to other age groups, which differs from these previous studies. However, voluntary ethanol consumption patterns vary with the species, (Green and Grahame, 2008) strain, (Yoneyama et al., 2008, Green and Grahame, 2008) and ethanol consumption paradigm utilized, (Green and Grahame, 2008) and prior studies noting increased ethanol intake in adolescent males used the drinking in the dark (DID) limited access paradigm (Moore et al., 2010) or utilized other mouse or rat strains. (Tambour et al., 2008, Morales et al., 2014) Similarly, prior studies in rats have found age-related increases in water consumption, (Cuitavi et al., 2024) though age-related decreases in water consumption have been observed in ethanol-exposed rats. (Garcia-Burgos et al., 2009)

Water-only mice showed a left-side preference which was more pronounced in males. Left side preference of bottles containing equivalent solutions has been noted in previous studies, which was hypothesized to indicate lateralization of consumption behavior. (Smutek et al., 2014) In the current study, the left bottle was closer to the cage wall, which could denote a form of thigmotaxis and/or neophobia of the sipper apparatus. Subsequent redesigns of sipper cage housing may place the left and right bottles on either side of the cage, such that both are next to the cage wall, to mitigate this behavior.

Binned analysis of automated home-cage sipper data revealed subtle differences in ethanol and water consumption across age, sex, and exposure groups, with interactions between age and sex in acrophase. Data binned every 60 minutes showed one major peak and one minor peak in consumption during the dark phase across most groups; a previous 2BC study utilizing home-cage lickometers similarly noted two peaks in ethanol consumption during the dark phase. (Caruso et al., 2021) Cosinor modeling of circadian drinking patterns showed minor phase advancement in ethanol consumption with age in females, but delay in males. Total fluid consumption (water + ethanol) acrophase was slightly earlier in 18-week females relative to other age groups, while 18-week males showed a mildly later acrophase than other age groups. In the water-only group, 6-week females showed a slightly delayed phase relative to other ages, while males showed slight phase delay with age. Direct comparison of circadian total fluid consumption data between exposure groups revealed a female-specific phase delay across age groups with ethanol exposure, as well as weaker amplitude that likely driven by adolescents.

Few-to-no studies examine circadian differences in ethanol consumption patterns by age and sex in rodents. However, one study found adolescent mice were resistant to chronodisruption with acute (intraperitoneal) ethanol exposure compared to adults. (Ruby et al., 2017) The current study presents voluntary ethanol consumption-related circadian rhythms and their potential phase disruption by age and sex in C57BL/6J mice that can be elucidated in further lickometer studies. Further studies should compare sex-specific circadian drinking patterns in adolescent, young adult, and mature adult drinking to aged mice, as alcohol consumption among adults over 65 has been increasing over the last 20 years and has also displayed a narrowing trend between males and females. (Keyes, 2023) Studying differences in ethanol consumption in advanced age will be critical given the United States population over 65 is expected to double by 2060. (2025)

This research has demonstrated the open-source automated home cage sipper devices developed by Godynyuk *et al*. (Godynyuk et al., 2019) are an effective means to further elucidate temporal ethanol consumption patterns by age and sex. Future research should elaborate on the current findings, including using these devices to compare ethanol drinking behavior in aged mice to younger age groups. Given the bidirectional relationship between circadian disruption and AUD (Hasler et al., 2015) and the sex- and age dependence of circadian rhythms (Ruby et al., 2017, Bellfy et al., 2025, Chang et al., 2026) and ethanol consumption (Ruby et al., 2017, Foo et al., 2023, Piekarski et al., 2022, Waldron et al., 2024, Collins et al., 1975, Melon et al., 2013, Mackowiak et al., 2025, Cuitavi et al., 2024, Garcia-Burgos et al., 2009) this avenue of study could help inform novel, therapeutics for AUD that account for differing chronobiology among age and sex.

## Data Availability

All sipper device housing files, data, and R code used in this current study are publicly available at https://github.com/rachelcrice/Homecage_Sippers.

