## Supplementary Figures for "Automated home-cage sipper devices reveal age and sex differences in ethanol consumption patterns"

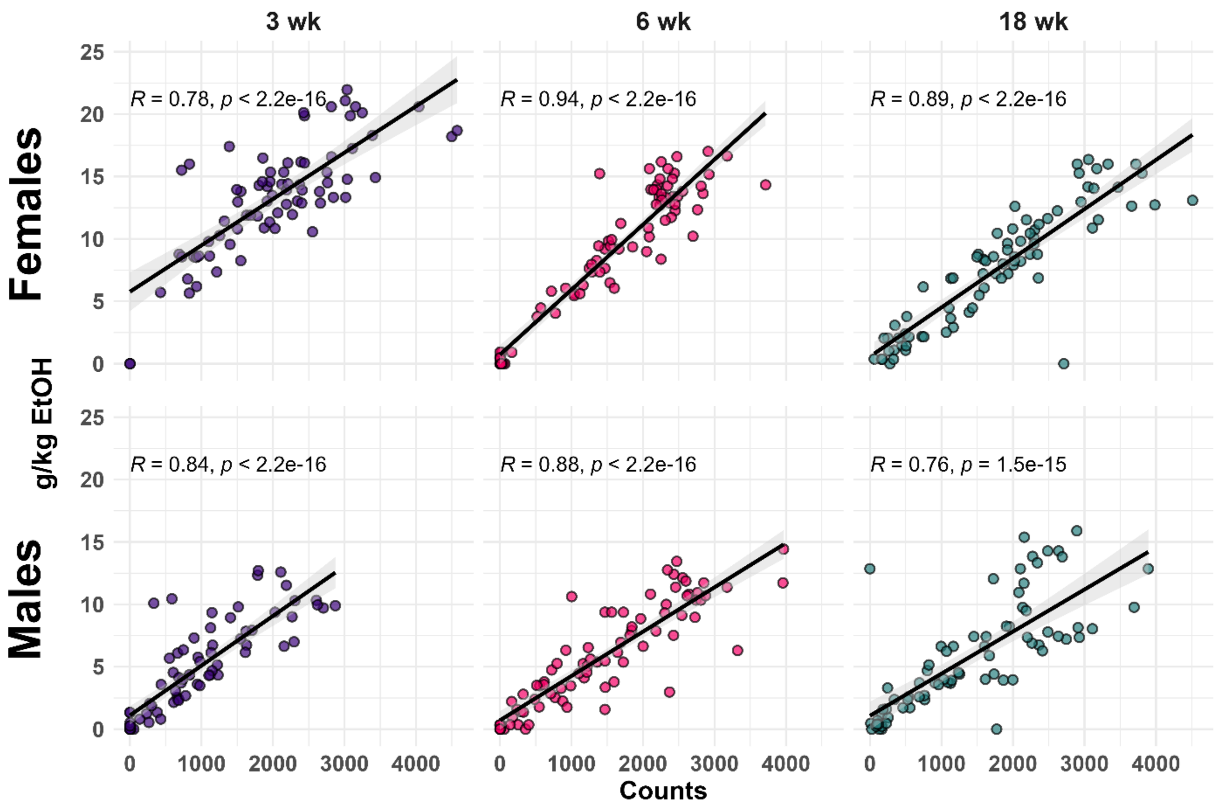


**Supplementary Figure S1. Correlations of manual 2BC ethanol consumption data versus sipper counts by age and sex.** The line of best fit is pictured for each group, with the 95% confidence interval shaded. All Pearson correlations were significant ($p < 0.001$) and strong ($R > 0.70$).

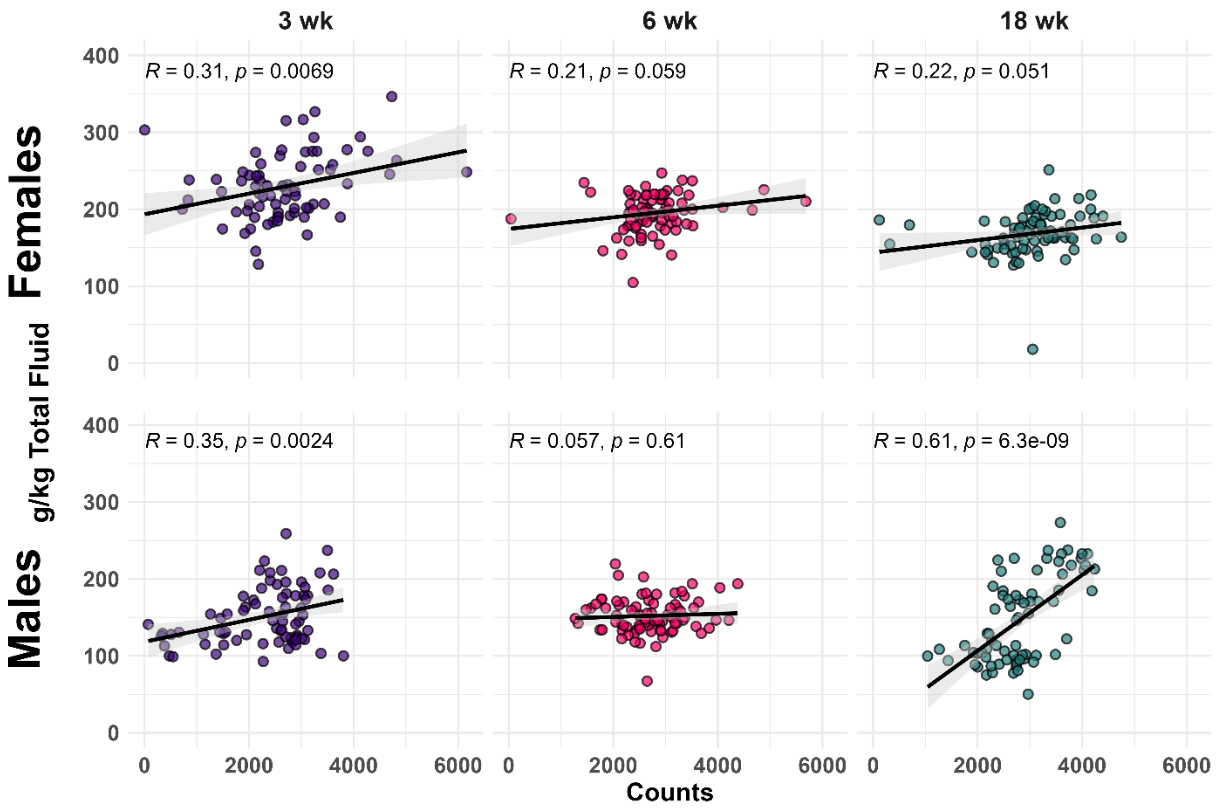


**Supplementary Figure S2. Correlations of ethanol group manual 2BC total fluid intake versus total sipper counts by age and sex.** The line of best fit is pictured for each group, with the 95% confidence interval shaded. Pearson correlations were significant for 3-week females and males ($p < 0.01$) and of weak strength ($R = 0.31$-$0.35$). Pearson correlation was also significant for 18-week males and of moderate strength ($R = 0.61$).

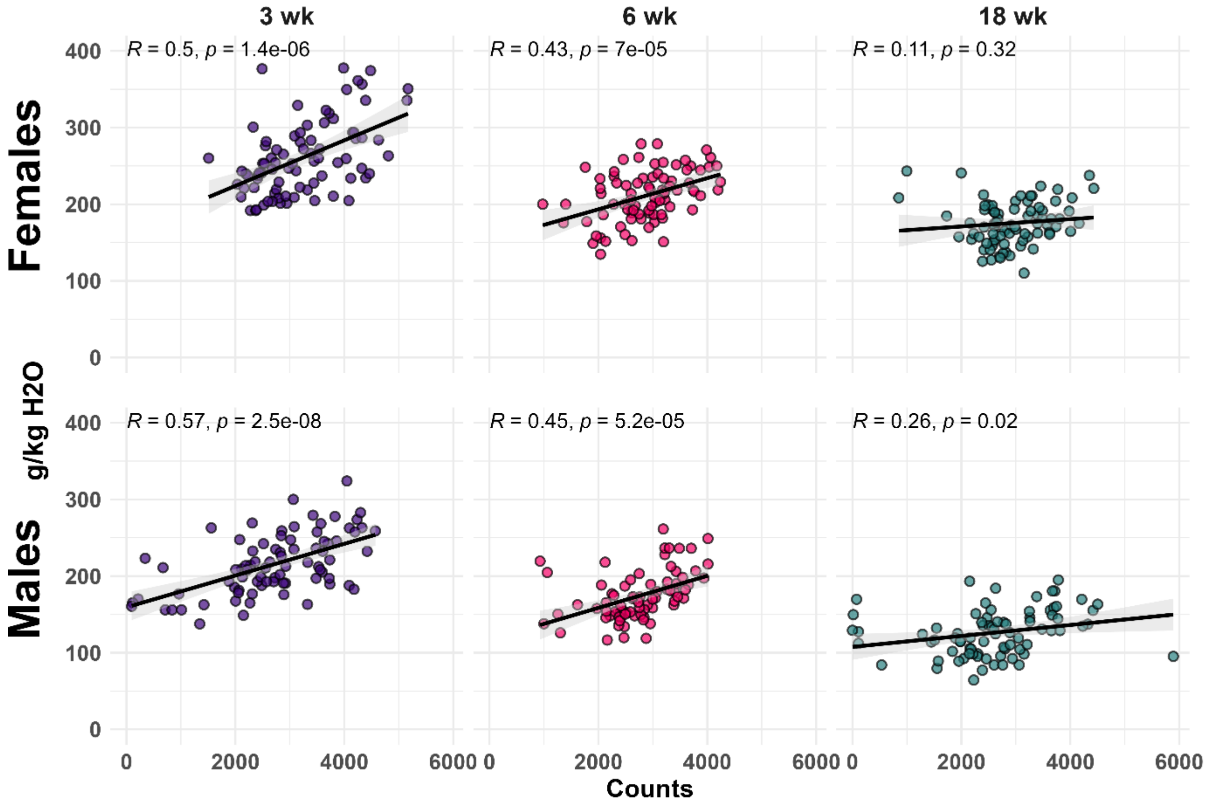


**Supplementary Figure S3. Correlations of ethanol-naïve group manual 2BC total water intake versus total sipper counts by age and sex.** The line of best fit is pictured for each group, with the 95% confidence interval shaded. All Pearson correlations except for 18-week females were significant and of weak-to-moderate strength.
